# Functional characterization of Neurofilament Light b splicing and *mis*balance in zebrafish

**DOI:** 10.1101/2020.04.27.064063

**Authors:** DL Demy, ML Campanari, R Munoz-Ruiz, HD Durham, BJ Gentil, E Kabashi

## Abstract

Neurofilaments (NFs), a major cytoskeletal component of motor neurons, play a key role in their differentiation, establishment and maintenance of their morphology and mechanical strength. The *de novo* assembly of these neuronal intermediate filaments requires the presence of the neurofilament light subunit, NEFL, which expression is reduced in motor neurons in Amyotrophic Lateral Sclerosis (ALS). This study used zebrafish as a model to characterize the NEFL homologue *neflb*, which encodes two different isoforms via splicing of the primary transcript (*neflbE4* and *neflbE3*). *In vivo* imaging showed that *neflb* is crucial for proper neuronal development, and that disrupting the balance between its two isoforms specifically affects NF assembly and motor axon growth, with resulting motor deficits. This equilibrium is also disrupted upon partial depletion of TDP-43, a RNA binding protein that is mislocalized into cytoplasmic inclusions in ALS. The study supports interaction of NEFL expression and splicing with TDP-43 in a common pathway, both biologically and pathogenetically.

## INTRODUCTION

A major cytoskeletal structure of the axon is composed by fibrillary networks of neurofilaments (NFs), which are expressed exclusively in mature neurons in both the central and peripheral nervous system [1] [2] [3]. NFs are formed by the assembly of three main proteins classified by their respective size: the *neurofilament light* (NEFL), *middle and high* protein (NEFM and NEFH) [4]. All NF proteins possess a conserved central alpha-helical rod domain, necessary for the dimer formation during the first step of assembly, and unique head and tail domains that protrude from the filament core [4]. The assembly properties of these factors depend on their stoichiometry [5], [6] and post-translational modifications [7], [8]. In fact, NEFM and NEFH cannot form long filaments by themselves, but each can co-assemble to form filaments in combination with NEFL both *in vitro* [9] and in cultured cells [10], [11]. NFs are especially abundant in motor neurons (MNs) where they play a key role in the organization of the neuronal cytoarchitecture and are involved in perikarya differentiation, protrusions (dendrites and axons) maturation and synaptic functions [12].

Any defect in NF assembly leads to dysfunction [13]. In particular, abnormal NF networks have been described in Charcot-Marie-Tooth disease (CMT) [14], [15] and Amyotrophic Lateral Sclerosis (ALS) [16].

In CMT, 30 mutations have been described in residues throughout NEFL [17] and cause a variable clinical phenotypes [18] from the complete absence of NEFL [19] to the production of aggregate-prone mutants [20], [21]. In particular, some of these mutants display altered phosphorylation patterns that suppress the filament assembly process, underlying the importance of this post-translational modification for NF assembly [21]. In ALS, the abnormal deposition of hyper-phosphorylated forms of NFs has been detected in MNs [22]–[24].The importance NEFL in ALS is supported by the following evidence. First, high levels of NEFL protein are detected in cerebrospinal fluid (CSF) and blood of all ALS cases [25] as early as 12 months before the onset of the disease [26]. Consequently, NEFL is considered the most relevant biomarker in ALS and its level is used in ALS prognosis as it strictly correlates with disease severity [27]–[29]. Second, perturbation of NEFL mRNA steady state in ALS spinal motor neurons could be a key pathogenic mechanism [30]. Indeed, disrupting the stoichiometry of NF subunits leads to NF aggregation reminiscent of ALS pathology [31]–[34]. The evidence of a pathological dysregulation of NEFL expression in ALS is strengthened by the existence of a direct interaction between NEFL mRNA 3’UTR region with SOD1, TDP-43, and RGNEF,, which are all genetic causes of ALS [35]–[39]. Importantly, the question of whether deficits in NEFL RNA regulation can induce motor neuron degeneration has not been tested *in vivo*. In order to address this hypothesis, we used zebrafish (*Danio rerio*), a vertebrate model used to easily study normal and pathological events in the nervous system [40]–[42]. A number of genetic models in zebrafish have been established for a range of neurodegenerative disorders, including ALS [43]–[49].

In this study, we examined the consequences of altering the expression of the NEFL orthologue (*neflb*) in zebrafish and established a direct link between *neflb* mRNA splicing modulations with an ALS-like phenotype (atrophy of motor axons and paralysis). We refined our understanding of this misbalance by ectopic expression of the two neflb isoforms, and also also explored *neflb* expression in a TDP-43 knockdown model.

## MATERIAL AND METHODS

### Zebrafish lines and microinjections

Wild-type and transgenic zebrafish embryos were raised at 28°C in embryo medium: 0.6 g/L aquarium salt (Instant Ocean, Blacksburg, VA) in reverse osmosis water containing 0.01 mg/L methylene blue. AB wild-type fish, and the transgenic lines Tg(Mnx1:eGFP) [50], Tg(elavl3:Gal4)zf349 - referred to as Tg(HuC:Gal4) - [51], Tg(Mnx1:Gal4) [52], Tg(5xUAS:RFP)^nkuasrfp1a^ - referred to as Tg(UAS:RFP) - [53] have been used in this study. Zebrafish husbandry was performed according to approved guidelines. All procedures for zebrafish experimentation were approved by the Institutional Ethics Committee at the Research Center of the ICM and by French and European legislation.

#### neflb-eGFP constructs/cloning

pUCminus containing N-terminally eGFP-tagged zebrafish cDNAs of both *neflb* splice variants (*neflbE3* and *neflbE4*) were purchased from Cliniscience. eGFP-*neflbE3* and eGFP-*neflbE4* were removed by restriction enzymes, and subcloned by ligation into pCS2 for ubiquitous expression in SW13 cells and p5e-10xUAS [54] for *in-vivo* expression in zebrafish motoneurons using the Tg(Mnx1:Gal4) trigger line. *Gene knock down using antisense oligomorpholino oligonucleotides (Mo):* Antisense Mo were designed complementary to bind ATG or splice junctions that would block transcription of the zebrafish targeted genes and synthesized from GeneTools (Philomath, USA). The sequence of the previously described *TDP43* Mo [43] is 5′-GTACATCTCGGCCATCTTTCCTCAG −3′. A splice-blocking antisense Mo against the *neflb* intron3-exon4 donor splice junction (*neflb*SV Mo) was synthesized (5′-CTCCCCTGTGAAAGAGTCAACAGAG −3′), and a standard control Mo - not binding anywhere in the zebrafish genome - was used to assess the specificity of the observed phenotypes (5’-CCTCTTACCTCAGTTACAATTTATA-3’).

1 nl of DNA constructs (final concentration 75ng/uL) and morpholino (0.2-0.6mM) solutions were microinjected in one-cell-stage embryos, using glass microcapillaries (Sutter Instrument) and a Picospritzer III (General Valve, Fairfield, NJ) pressure ejector.

### *In vivo* live imaging and timelapse

For live confocal fluorescence imaging, embryos were mounted in 1% low-melting-point agarose (16520050; Invitrogen) dissolved in embryo water, and supplemented with 0.16 mg/ml tricaine (A-5040; Sigma). After solidification, embryo medium with 0.16 mg/ml tricaine solution was added in order to keep embryos hydrated and anesthetized during experiments. Thereafter, images were captured at the selected times on an inverted Leica SP8 set-up allowing multiple point acquisition, so as to image mutants and their siblings in parallel. Time-lapse acquisitions were captured with a Spinning Disk system (Andor technology, UK; Leica Microsystems, Germany), a DMI8 inverted stand (Leica Microsystems, Germany), a CSU-X head (Yokogawa, Japan) and a QE-180 camera (Hamamatsu, Japan), with a 20x objective (NA0.5). Image stacks were processed with LAS software to generate maximum intensity projections or were exported into ImageJ or Imaris software (Bitplane) for 3D analysis of motor axons projections.

### Imaris filament tracer

For each embryo, a region of interest containing 4 somitic nerve fascicles (somites 13 to 16) was manually defined in 3D, then neurofilaments were analyzed using the Filament tracer plugin of the Imaris software (Bitplane). Filament length, filament total volume, filament full branch depth and filament full branch level values were exported to Excel (Microsoft) and Prism (GraphPad).

### Acridine orange staining

Apoptotic cells were revealed in 2-days-old live embryos by adding the Acridine Orange vital dye (N-4638; Sigma) to the embryo water at a final concentration of 5 mg/ml. Embryos were then incubated in the dark at 28°C for 1.5–2 h, rinsed, anesthetized and observed under a fluorescent stereomicroscope (Olympus, Japan).

### Touch Evoked Escape Response (TEER)

At 48 hpf, zebrafish embryos were tested for motor behavior under a stereomiscroscope (Zeiss, Germany) using the TEER experiment. Morphologically normal zebrafish from each experimental condition were touched lightly at the level of the tail with a tip, and their responses to the stimulus were recorded with a Grasshopper 2 Camera (Point Grey Research) at 30 Hz. The videos were then analyzed using the Manual Tracking plugin of ImageJ software, and the swim duration, swim distance, and maximum swim velocity of each embryo were calculated as previously described [44].

### FACS

Embryo dissociation was performed as previously described [55]. Sorted cells were collected in phosphate-saline buffer (PBS) and RNA was extracted using Trizol reagent (T9424, Sigma). The extraction, RT and PCR were performed on three biological replicates of 20.000 cells each.

### Cell culture

The human cells SH-SY5Y were grown in D-MEM (Dulbecco’s Modified Eagle medium; Gibco®, Life technologies Paisley, UK) supplemented with 10% fetal bovine serum and 1% of penicillin/streptomycin solution (100 U/mL). Cells were seeded at a density of 8×105 cells on 35 mm tissue culture dishes and were transfected the following day with TDP43 siRNA using Lipofectamine™ 2000 (Invitrogen™, Life technologies Paisley, UK) according to the manufacturer’s instructions. After 48 hours of transfection, cells were washed with PBS and resuspended in 130 µL ice-cold extraction buffer: 50 mM Tris-HCl, pH 7.4 / 500 mM NaCl / 5 mM EDTA / 1% (w/v) Nonidet P-40 / 0.5% (w/v) Triton X-100 supplemented with a cocktail of protease inhibitors. Cell lysates were then sonicated and centrifuged at 14000g at 4 ºC for 20 minutes. The supernatants were collected and frozen at −80ºC until biochemical analysis.

SW13vim-cells, which lack endogenous intermediate filaments, were cultured in Dulbecco’s Modified Essential Medium with 5% fetal bovine serum (FBS). Cells were transfected with Lipofectamine 2000 in Optimem medium (Invitrogen, Carlsbad, CA, USA) according to the manufacturer’s instructions using plasmids encoding, EGFP-*neflbE4*, EGFP-*neflbE3*, mouse NEFM (identified by the clone NN18, Sigma–Aldrich, 1:1000) and NEFH (identified by the clone N52, Sigma–Aldrich, 1:1000), as previously described [56].

### Reverse Transcription and PCR

Total RNA - from zebrafish embryos, FACS-sorted cells or cell cultures - was extracted using the TRIreagent (T9424, Sigma). RNA was quantified using the Nanodrop 8000 (Thermo Scientific) and its quality was checked using the 2100 Bioanalyzer (Agilent Technologies). cDNA was synthesized from 1ug of RNA using Transcriptor Universal cDNA Master Mix (Roche). PCR experiments were performed to assess the expression of genes using primer pairs amplifying the sequences of interest **(Supplementary table 1)**.

### Western Blot

Embryos at the stage of interest were deeply anesthetized, and collected into 1.5mL microcentrifuge tubes. Water and anesthetic were completely removed and replaced by 200uL of RIPA solution supplemented with protease inhibitors. Tissue dissociation was performed by ultrasound, and solubilized proteins were quantified with the Pierce BCA protein assay (Thermofisher).

Proteins were denatured for 7 min at 98°C, separated on 4–12% bis-Tris Gel (NuPAGE), transferred on nitrocellulose membranes (Whatman Protran), blocked with 5% BSA in Tris-buffered saline (TBS) buffer, incubated with Tubulin (T5168, Sigma) or TDP43 (3449S, Cell Signaling) antibodies diluted in TBS plus 5% BSA, washed three times, and incubated with anti-rabbit or anti-mouse dye-conjugated antibodies (1:3,000, Cell Signaling) for 1 h in TBS, followed by washing and the signal was detected using the ODYSSEY® CLx. The intensity of bands was measured by the open source image processing program ImageJ.

### Statistical analysis

Data were plotted and analyzed using the Excel software (Microsoft, USA). To evaluate differences between means of non-Gaussian data, they were analyzed with a Mann-Whitney-Wilcoxon (for comparison of two groups), combined with a Kruskal-Wallis test (for multiple comparison). When appropriate, distributions were normalized by log, and an analysis of variance (Two-way ANOVA) was performed (for experiments with two variables). P<0.05 was considered statistically significant (symbols: ****P<0.0001; ***P<0.001; **P<0.01; *P<0.05). Statistical analyses and graphic representations were done using Prism software. Additional details of each test are described in related figures legend.

## RESULTS

### The zebrafish homolog *neflb* encodes two splice variants, *neflbE3* and *neflbE4*, differently expressed during the development

The zebrafish genome harbors a predicted NEFL homolog, *neflb*, based on its sequence similarity with the human gene (Ensembl ID: ENSDARG00000012426). It displays about 60% of nucleotide sequence identity with its mammalian homolog; the exon/intron structure has been mostly conserved through evolution, and two splice variants have been predicted for this gene (one with three exons - *neflbE3* - and one with four exons – *neflbE4*) in zebrafish (Figure 1A), as well as in humans (Ensembl ID: ENSG00000277586). In order to study *neflb* expression through gastrulation and organogenesis, we extracted RNA from zebrafish embryos at 3, 6, 24, 48, 72 and 96 hours post fertilization (hpf), and designed primers so to reveal both splice variants and differentiate them by their size. As shown in Figure 1B, *neflb* is expressed in zebrafish at all tested stages, and both expected splice variants were detected. *neflbE3* (upper PCR band), was detected only from 3 hpf up to 24 hpf, while the other variant, *neflbE4* (lower PCR band), was expressed from 24 hpf. This temporal shift in expression correlates with neurogenesis and with the stage at which *neflb* mRNA, detected by *in situ* hybridization, stopped being expressed ubiquitously, and started being expressed specifically in developing neurons [57]. To determine whether the *neflbE4* isoform was indeed specifically expressed in neuronal cells, we used Tg(HuC:Gal4/UAS:RFP) embryos. In this double transgenic line, all post-mitotic neurons (HuC positive) express RFP (Figure 1C, *i-i”*); after FACS-sorting of RFP+ neurons from RFP-non neuronal cells (Figure 1D), RNA was isolated and PCR was performed to detect mRNA for genes of interest with β-actin as a control. As shown in Figure 1E, both cell pools express similar levels of β-actin mRNA, whereas RFP mRNA was only expressed in the neuronal pool. *neflbE4* mRNA was detected only in post-mitotic neurons.

**Figure 1.**
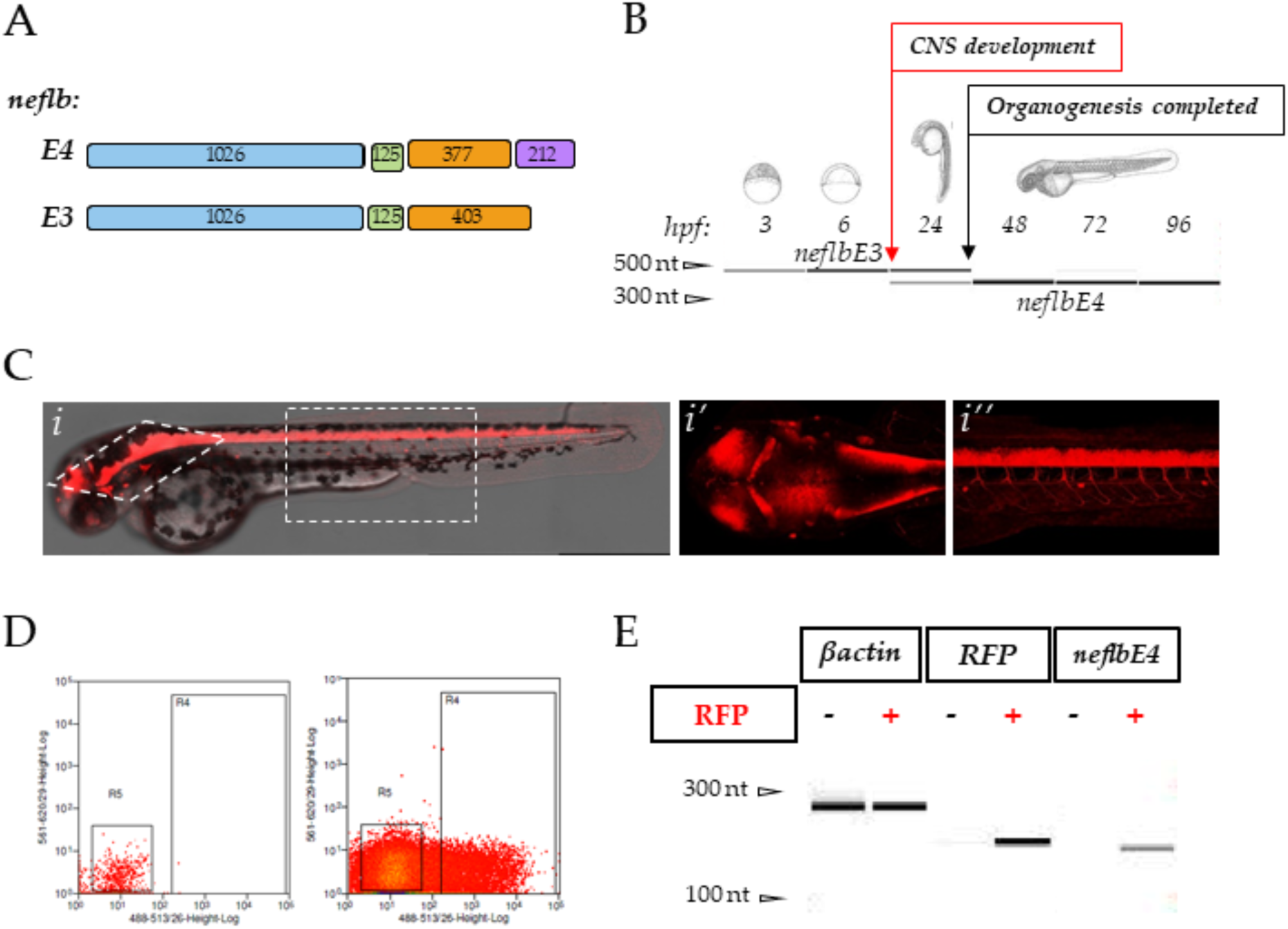
*neflbE3* and *neflbE4* expression in zebrafish. **A**, schematic of the zebrafish exon structures of the predicted *neflb* splice variants. Length (bp) is indicated on each exon**. B**, *neflb* is expressed at all embryonic and larval stages in zebrafish, with a splicing shift from *neflb3E* (upper PCR band) to *neflb4E* (lower PCR band) occurring during CNS development - revealed by a change in the amplicon size. **C** (*i-i’’*), sagittal and dorsal vision of Tg(HuC:Gal4/UAS:RFP) zebrafish embryos at 48hpf. This transgenic line express the red fluorescent protein (RFP) in post-mitotic neurons. **D**, post-mitotic neurons (RFP+) and non-neuronal cells (RFP-) were isolated by FACS, and expression of β-actin, RFP and *neflb4E* was tested by PCR in both pools (**E**).

### *neflbE3/E4* misbalance results in a strong and specific motor phenotype

In order to study the role of *neflb* splicing and the importance of *neflbE4* in CNS development, we designed an antisense morpholino oligonucleotide (Mo) targeting the splice acceptor junction between intron 3 and exon 4 of the *neflb* gene (*neflb* SV Mo) (Figure 2A). This Mo was predicted to prevent the excision of intron 3, driving the splicing balance towards the generation of the *neflbE3* isoform.

**Figure 2.**
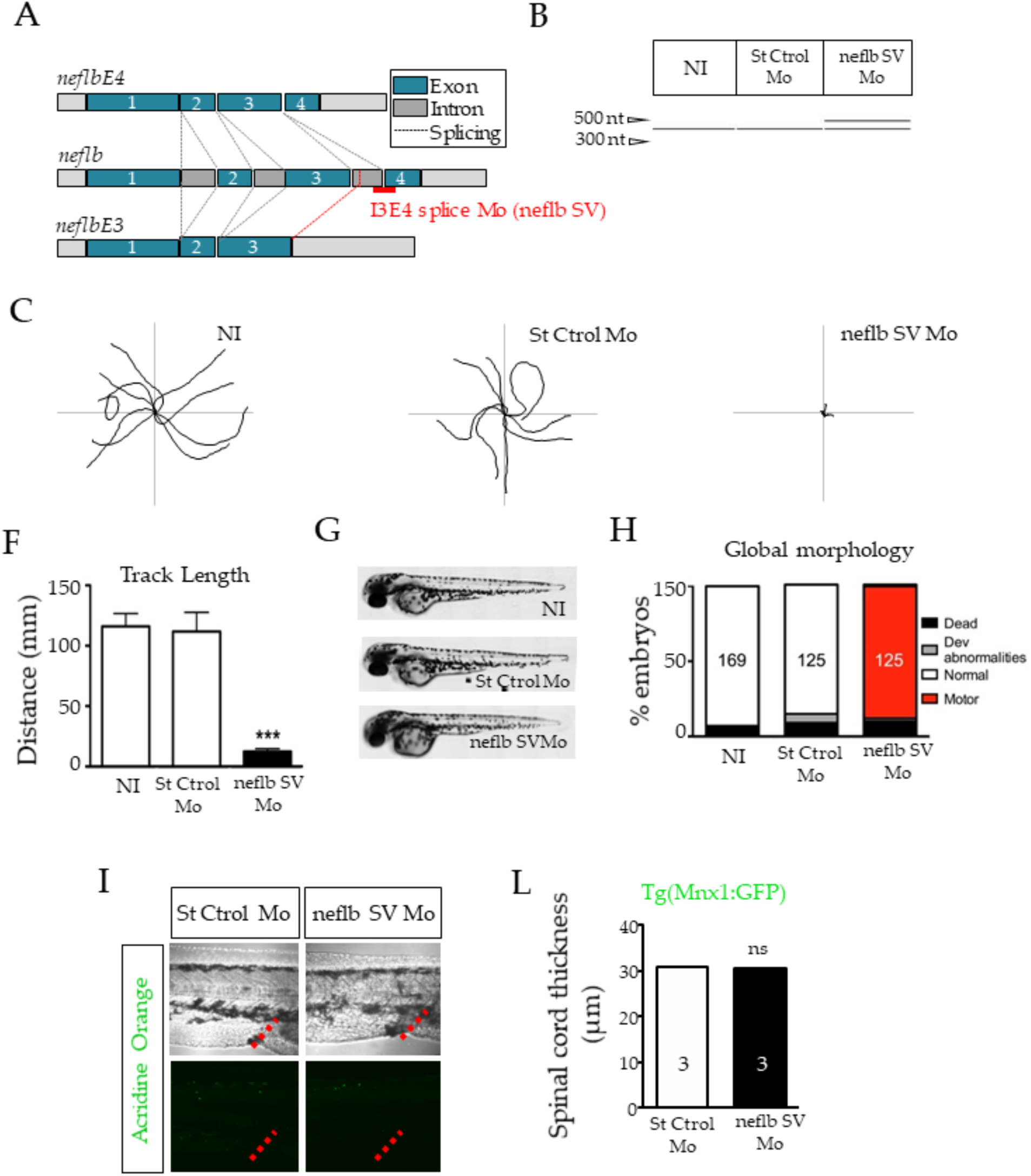
*neflbE3/E4* misbalance results in a strong and specific motor phenotype. **A**, position of the splicing variant (SV) antisense oligomorpholino (neflb SV Mo, in red) targeting the decisive I3E4 splice junction in order to inhibit the developmental shift from *neflb3E* to *neflb4E*. **B**, St Ctrol Mo and neflb SV Mo were injected at the dose of 0.2 mM. The I3E4 morpholino was efficient and caused the persistence of *neflb3E* (upper band) expression at 48 hpf, when only *neflb4E* (lower band) was expressed in controls at this stage. **C**, trajectories of 9 representative zebrafish embryos per condition during Touch Evoked Escape Response at 48hpf. Non-injected (NI) and control injected embryos (St Ctrol) swam to the edges of a Petri dish in reaction to a light touch; neflb SV morphants were unable to move away from the center of the dish. **F**, quantification of the track length shows a reduction by 90% of the swimming distance in neflb SV Mo injected fish compared to the controls. **G**, neflb SV Mo injected embryos develop without any major developmental abnormality. **H**, bar graph of the phenotypes distribution after the TEER analysis at 48 hpf. Percentages of zebrafish with motor deficits (motor) are increased after neflb SV Mo injection. Percentages of normal developed embryos in NI and St Ctrl Mo conditions are comparable. **I**, Acridine orange staining (green), which is a metachromatic intercalator sensitive to DNA conformation, is used in this study to detect apoptosis. No higher amount of Acridine Orange positive cells was detected in the spinal cord of neflb SV morphants (right panel) compared to the controls (left panel). The urogenital opening (red dotted lines) has been used as geographical reference for measures. **L**, spinal cord thickness measured at 48hpf. ***P<0.001; ns, non-significant.

As expected, at 48hpf only *neflbE4* (Figure 2B, lower band) was detected in control uninjected embryos, as well as in embryos injected with a standard control Mo (St Ctrol Mo). In contrast, in embryos injected with *neflb* SV Mo, an upper band appeared representing the splicing variant *neflbE3*, which expression is maintained beyond 24 hpf.

We then characterized the motor phenotype of each condition at 48 hpf by Touch Evoked Escape Response (TEER). Following a light touch, both uninjected and St Ctrol Mo injected embryos escaped and swam away from the center of the Petri dish all the way to the edges of the plate, whereas embryos injected with *neflb* SV Mo did not (Figure 2C), swimming distance being reduced by 89% (Figure 2F).

Although no developmental abnormalities were observed among conditions (Figure 2G), *neflb* SV Mo injection resulted in a very strong and specific motor phenotype (88% of embryos) characterized by the inability to swim (Figure 2H). As no differences were observed between uninjected and St Ctrol embryos, only the later were used further in this study.

To determine whether the motor deficit observed was associated with cell death, Acridine Orange vital staining was used to detect apoptotic neurons. No major signal was detected in *neflb* SV Mo embryos compared to controls (Figure 2I). There were also no significant differences in spinal cord thickness among the groups (Figure 2L). Thus, we concluded that the abnormal expression of *neflbE3* isoform doesn’t affect neurons development.

### The motor phenotype associated with the alteration of *neflbE3* expression correlates with atrophy of motor axons

Previous work has shown a direct link between motor behavior deficits and disorganization of motor neuron morphology [44], [58]–[60]. Thus, we examined the somitic axonal projections of the motor neurons. Axonal projections from motor neurons exit the spinal cord grouped as one nerve fascicle per somite, then grow along the somitic muscle all the way to its most ventral part and then back up around it, while branching and connecting with muscle fibers. Mirror innervation of the dorsal part of the somitic muscle is achieved through the same process.

As shown in Figure 3A *i*, all somitic muscles appeared properly innervated in control embryos. However, neflb SV morphants displayed shortened and disorganized motor neuron nerve fascicles with major branching abnormalities (Figure 3A *ii*). Filament tracer plugin of the Imaris software was used to model the nerve fascicles in 3D and analyze the morphological defects (Figure 3A *i’* and *ii’*). In neflb SV morphants, the somitic nerve fascicle main length was decreased by 46% as compared to controls (Figure 3C). Furthermore, axons were less branched (Figure 3D) and innervating a reduced area on muscle (Figure 3E), confirming the strong and specific phenotype of the neflb SV morphants.

**Figure 3.**
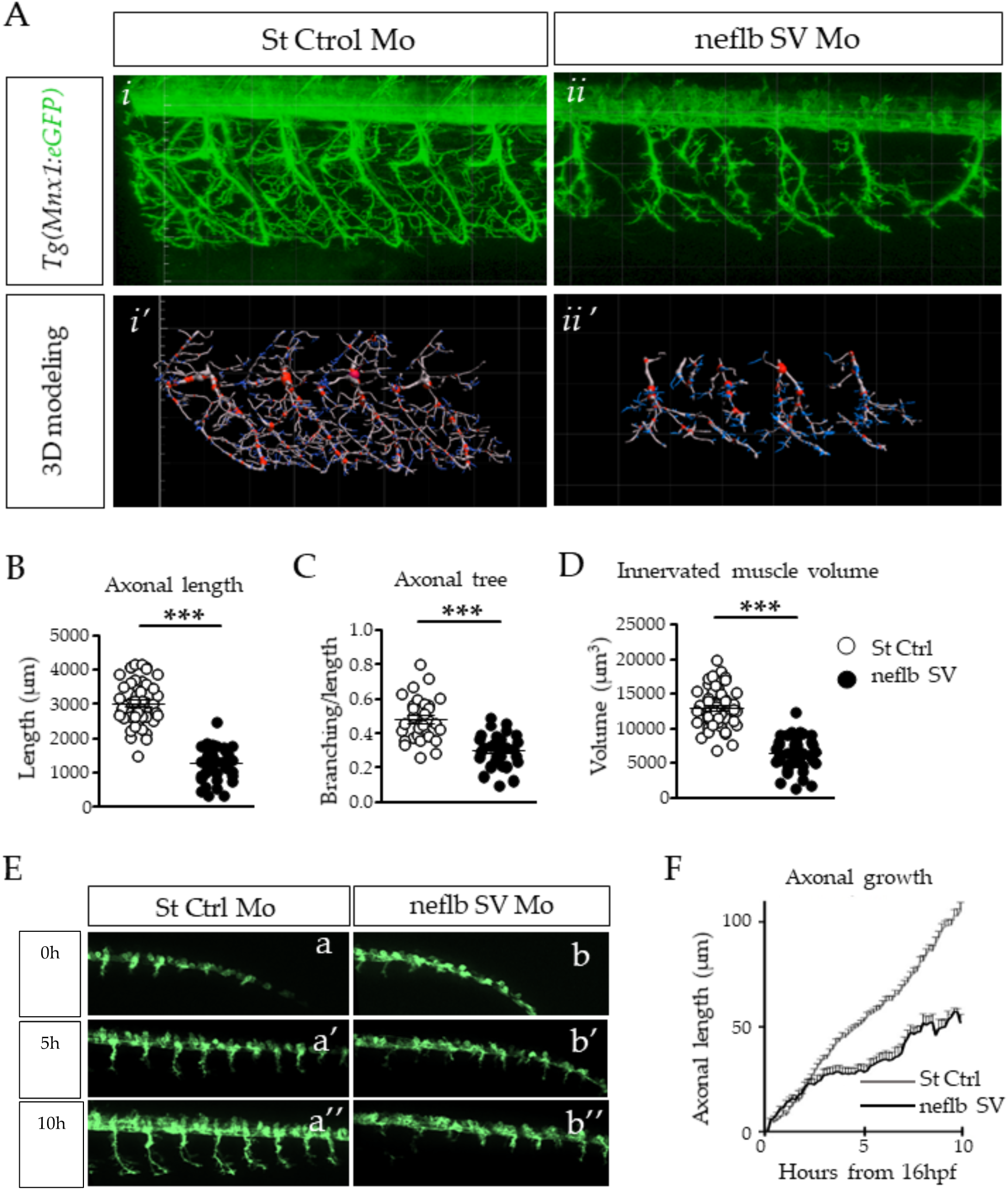
Axonal atrophy in motor neurons of neflb SV embryos. **A**, *In vivo* observation of somitic nerve fascicles in 48hpf Hb9:eGFP zebrafish embryos injected with a control Mo (*i*) or with neflb SV Mo (*ii*). Hb9:eGFP line express GFP (green) in motor neurons under the specific promoter HB9. After neflb SV Mo injection axonal projections appeared shorter and less regularly distributed and branched than controls. *i’* and *ii’*, 3D modeling of motor neurons morphology using the Imaris software. This reconstruction permitted the quantitative analysis of the nerve fascicles features**: B**, the axonal length; **C**, the axonal branching; **D**, the volume of the innervated muscle. All measurements were reduced in neflb SV morphants (black circles) respect with the controls (white circles). **E**, i*n vivo* time-lapse imaging of *Tg(Mnx1:eGFP)* zebrafish embryos injected with a control morpholino (a-a’’) or with the neflb SV morpholino (b-b’’) was performed during 10 hours starting from 16 hpf. **F**, quantification of motor neurons axonal length during time. hpf, hour post fertilization. ***P<0.001.

To refine our understanding of this phenotype, we performed *in vivo* time-lapse imaging over the first ten hours (starting from 16hpf) of muscle innervation by motor axons, in control and nefl SV morphants in parallel. As shown in **Supplementary Movie 1** and in Figure 3E, motor axonal projections in *nefl* SV morphants appear to initiate at the same developmental stage as in the controls (**Figure E a- b’**), and they sprout in the right direction, ventrally towards the somitic muscles. However, as shown in **Supplementary Movie 1**, motor axons constantly grew back and forth, and never reached normal length (Figure 3E a’’-b’’, Figure 3F).

### *neflbE4* participates in motor neuron axonal growth, whereas *neflbE3* fails to polymerize normally and forms aggregates

In order to assess the assembly properties of *neflbE4* and *neflbE3 in vivo*, we generated both UAS:eGFP-*neflbE4* and UAS:eGFP-*neflbE3* constructs. These constructs were separately injected into Tg(Mnx1:Gal4/UAS:RFP) zebrafish embryos, in order to target the expression of the two isoforms in motor neurons.

At 48hpf, eGFP-*neflbE4* positive motor neurons have the same morphology as in eGFP+ controls **(Supplementary Figure 1A).** The eGFP-*neflbE4* protein is homogenously distributed throughout cell bodies, axons and dendrites, (Figure 4A *i-ii*) and is abundant in all axonal ramifications (Figure 4A *a-c*, **dotted circles)** and NMJ boutons (Figure 4A *a-c*, **arrows).** In contrast, the expression of the eGFP-*neflbE3* isoform induced changes in motor neuron morphology, with loss of dendrite extensions. In addition, **the** eGFP-*neflbE3* isoform formed cytoplasmic bundles (Figure 4B *i*) and aggregates (Figure 4B *ii*) in almost all motor neurons expressing the construct **(Supplementary Figure 1B)** and impaired the formation of axonal ramifications and NMJ (Figure 4B *a-c*).

**Figure 4.**
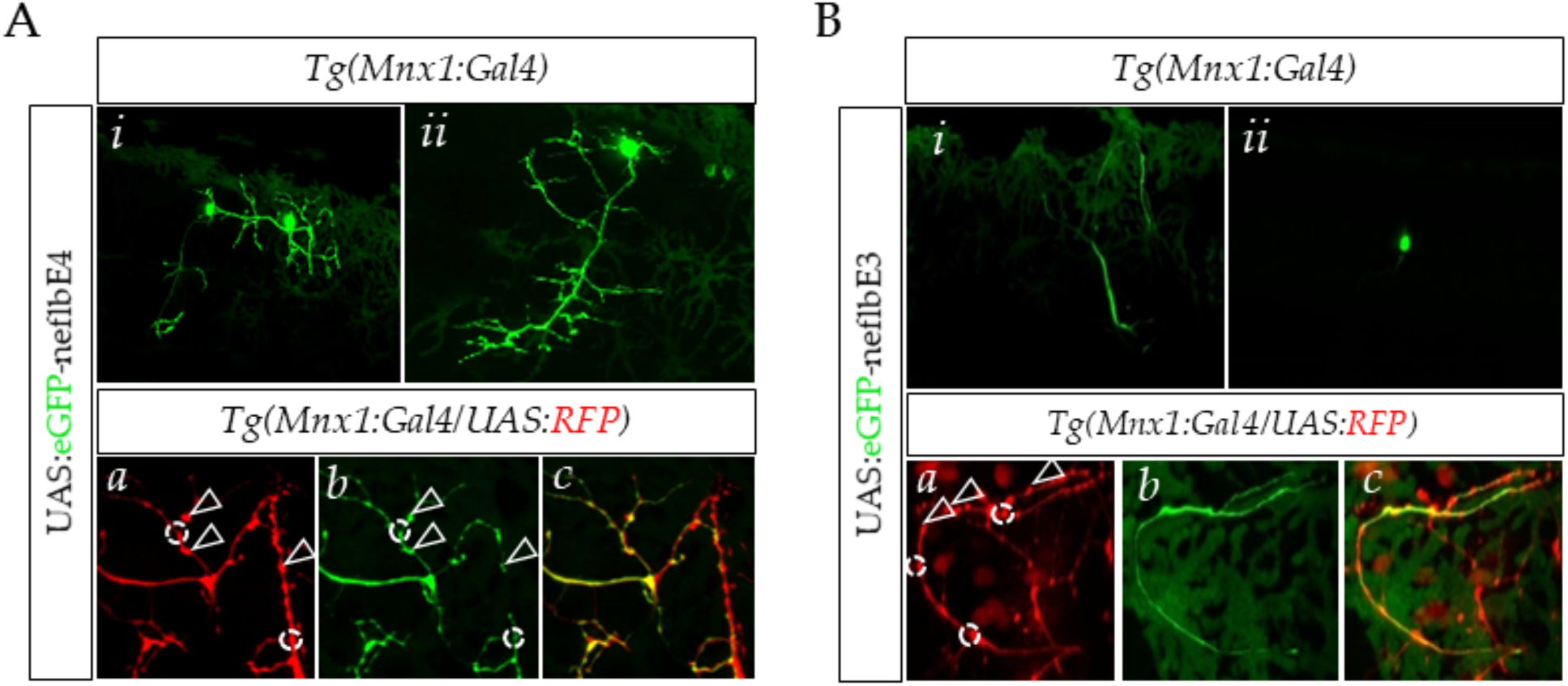
neflbE4 and neflbE3 distribution within motor neurons. **A** *(i-ii*), *in vivo* observation of single motor neurons expressing eGFP-neflbE4. Motor neurons expressing eGFP-neflb4E extend long and ramified axons. *(a-c*), eGFP-neflbE4 is particularly abundant in specific area representing the axonal branching (white dotted circles) and NMJ buttons (white arrows). **B** *(i-ii*), single motor neurons expressing eGFP-neflb3E. eGFP-neflbE3 mostly aggregated or assembled into long, massive, unramified bundle. *(a-c*), eGFP-neflbE3 was reduced or absent in axonal ramifications sites and NMJ buttons.

### *neflb* splice variants have different assembly properties in SW13^vim-^cells

To evaluate assembly properties, the constructs were expressed in SW13^vim-^cells, which lack endogenous intermediate filament proteins. In mammalian cells, NEFL dimerizes with NEFM and NEFH and is a core protein necessary for the assembly of NF proteins into filamentous structures [61]. In order to determine the assembly competency of *neflb* variants, eGFP-*neflbE4* or eGFP-*neflbE3* were expressed alone or in combination with mouse NEFM or NEFH (Figure 5A).

**Figure 5.**
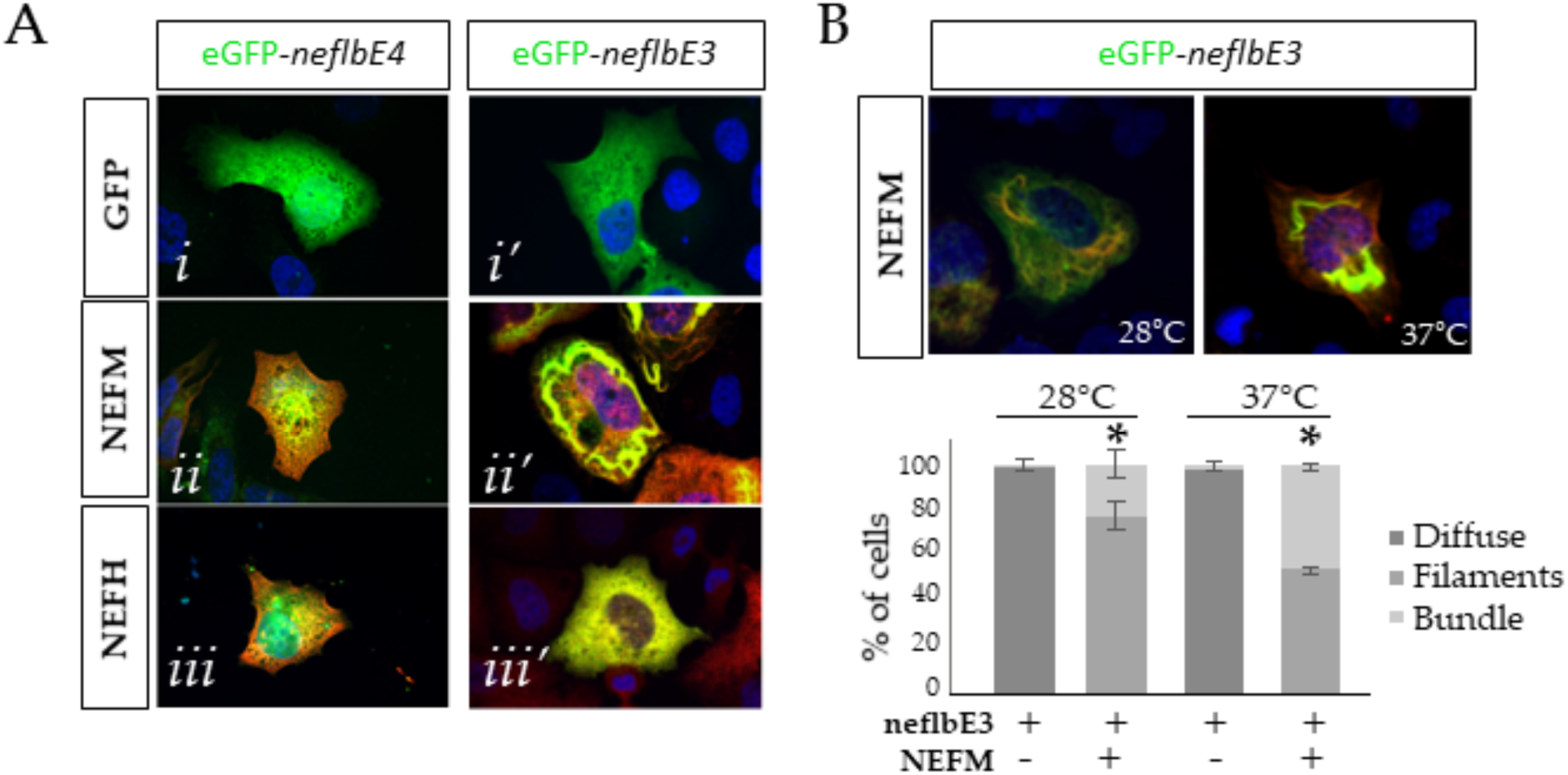
*neflb* splicing variants assembly properties in SW13vim-cells. **A**, eGFP-neflbE4 (*i*) and eGFP-neflbE3 (*i’*) were expressed in SW13vim-cells alone or together with mouse NEFM (*ii-ii’*, red) or NEFH (*iii-iii’*, red). The assembly pattern was characterized at 48h post transfection at 37°C. **B**, eGFP-neflbE3 and NEFM were co-transfected and bundle formation was look at 37°C and 28°C. *Bottom panel*, Bar graph showing the percentage of cells carrying filaments, bundles or diffuse labeling. At 28°C, the proportion of cells carrying bundles made of neflb3E and NEFM was less than the proportion of cells carrying bundles at 37°C. *p< 0.05, (n=3 cultures, 60 to 100 cells per coverslip) one-way ANOVA Tukey’s HSD post-hoc analysis.

Neither *neflbE4* nor *neflbE3* assembled into filamentous structures on their own (Figure 5A *i*-*i’*). Although eGFP-*neflbE4* and eGFP-*neflbE3* partially co-localized with NEFM suggesting an interaction, only eGFP-*neflbE3* could form higher order structures (filaments and bundles) with NEFM (Figure 5A *ii’*). Finally, neither of them formed filaments in presence of NEFH (Figure 5A *iii*-*iii’*).

Because zebrafish are poikilothermic organisms living at 28°C, *neflb* could have different assembly properties according to temperature, as previously shown for the lamprey Nefl [62]. Therefore, assembly of *neflbE3* with NEFM was verified not only at 37°C, but also at 28°C (Figure 5B). At 28°C, *neflbE3* was still able to form bundles of filament with NEFM, although less efficiently than at 37°C. Thus, the basic mechanisms of assembly and bundling of *neflbE3* are operant at both temperatures.

### TDP-43 regulates *neflb* splicing in zebrafish

Because of the role of NFs and TDP-43 in ALS pathology, we aimed to determine if *neflb* splicing occurs in a well characterized zebrafish model of ALS [43], where TDP-43 is depleted by morpholino injection. The knockdown efficiency of TDP-43 expression was measured by Western blot analysis (**Supplementary Figure 3A**) and zebrafish showed significant reduction in swimming parameters at 48hpf TDP-43 KD (**Supplementary Figure 3B**) as well as decreased somitic muscle innervation (**Supplementary Figure 3C**).

Interestingly, in the absence of TDP-43, *neflb* intron 3 was retained (Figure 6A), suggesting an important role for TDP-43 in *neflb* splicing, with consequent impact on regulation of NF expression and maintenance of NF protein stoichiometry, affecting motor neuron morphology and function.

**Figure 6.**
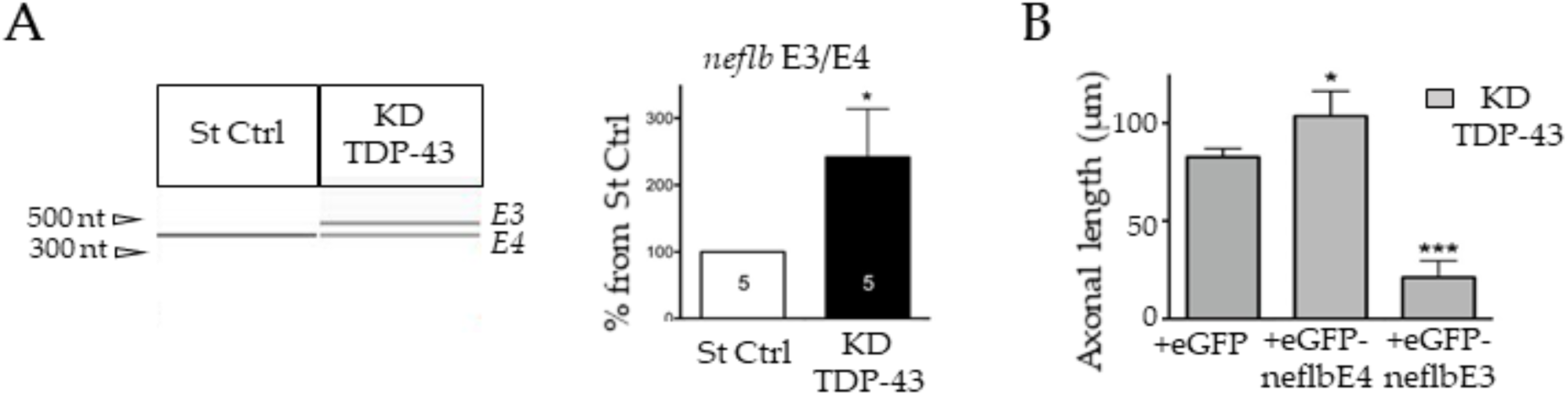
TDP-43 regulates *neflb* splicing in zebrafish. **A** *left*, PCR analysis on RNA extracted from St Ctrl embryos and fish KD for TDP-43, using specific primers for both isoforms *neflbE4* and *neflbE3*. *Right*, quantification of *neflbE3* and *neflbE4* ratio. **B**, axonal length measurement at 48hpf after over expression of eGFP-*neflbE4* and eGFP*-neflbE3* constructs in KD TDP-43 embryos. *neflbE4* significantly rescued axon length (middle histo bar), whereas *neflbE3* significantly aggravated the KD TDP-43 phenotype (right histo bar). *P<0.05, ***P<0.001.

To verify this hypothesis, we injected the previously described eGFP-*neflbE4* and eGFP-*neflbE3* constructs into TDP-43 KD embryos expressing Tg(Mnx1:Gal4) and compared motor neuron axonal length among conditions (Figure 6B). Interestingly, the reduced axonal length of motor neurons of TDP-43 KD embryos (eGFP) was rescued by eGFP-*neflbE4* injection, showing that *neflb* splicing is a key event in the pathogenic cascade leading to abnormal motor neuron architecture. Importantly, eGFP-*neflbE3* isoform expression exacerbated such deficit showing that the proper splicing of *neflb* is key for motor neuron.

## DISCUSSION

### General Results

Post transcriptional regulation of *NEFL* can have a wide impact on the NF network. However, the relevance of *NEFL* mRNA metabolism in ALS pathogenesis is unclear. In this study, we addressed the function of zebrafish *neflb* processing in regulation in motor neuron development and pathogenic consequences of dysregulation. Only recently, the *NEFL* orthologue *neflb* has been characterized in zebrafish, and shown to be expressed in neurons [57]. Here, we described the existence of two mRNA splice variants of this gene that we called *neflbE4* and *neflbE3*, which are differently expressed during zebrafish development. *neflbE3* is the first isoform to be produced ubiquitously before being completely replaced by *neflbE4*, which is a neuron-specific isoform. Interestingly, when *neflbE3* isoform expression was forced to persist beyond its developmental window, the fish displayed a strongly compromised swimming motor behavior, which correlated with aberrant motor axons growth and ramification. Importantly, these affected motor neurons present an abnormal assembly of *neflbE3* protein consistent with the formation of abnormal bundles and aggregates when expressed together with NEFM in SW13^vim^, human cell line lacking intermediate filaments commonly used to study *de novo* NF assembly.

**NF stoichiometry:** The requirement for NEFL in the assembly of the other subunits, NEFM and NEFH, has been widely described [6], [63], [64]. Thus, Nefl KO mice exhibit motor neurons with no NFs and axons of smaller caliber [65], as well as functional consequences [66]. Furthermore *NEFL* mRNA is reduced in ALS motor neurons in autopsy specimens and is associated with altered NF triplet protein stoichiometry and neurofilamentous aggregation [30].

The relevance of disruption of NF stoichiometry has also been implicated in conditions of NEFL overexpression. Transgenic overexpression of Nefl in mice was sufficient to generate NF accumulation in motor neuronal perikarya and proximal axons, accompanied by axonal and dendritic degeneration [32].

On the contrary, the additional expression of NEFL was beneficial in a transgenic mouse overexpressing NEFH [67], reducing NF accumulation and axonal transport defects in a dose-dependent manner. Taken together, these results reinforce the importance of the precise stoichiometric abundance of NEFL for proper NF assembly.

In our zebrafish model, the re-expression of the early isoform of *neflb* (*neflb3E*) at later stages induced motor and morphological phenotypes reminiscent of what is observed in transgenic mice and ALS patients. The ensemble of our results confirm the importance of NFs stoichiometry for polymerization and the existence of splicing mechanisms in regulation of NF biology in zebrafish.

### TDP-43 and splicing

mRNA *mis-*splicing is a common pathological mechanism widely recognized in ALS. In particular, the effects of TDP-43 on RNA metabolism have been extensively studied due to the pivotal role that this protein plays in ALS physiopathology [68]–[72]. TDP-43 mutations are known to induce ALS clinical traits through mRNA *mis-*splicing in patients [73], [74]. Depletion of TDP-43 in cell culture can cause the abnormal splicing of cryptic exons and lead to toxicity [75]. Also, mice expressing TDP-43 mutated in its low-complexity domain suffered from neurodegeneration that correlated with the consistent skipping of constitutive exons normally spliced by wild-type TDP-43 (named skiptic exons) [72], [76]. In 2007, the direct stability-mediated effect of TDP-43 on *NEFL* mRNA was described [77]. In this study, we explored the influence of TDP-43 on *neflb* gene expression in zebrafish using an ALS model generated by *Tardbp* KD [43]. TDP-43 loss of function induced retention of the intron 3 of the zebrafish *neflb* gene, with the consequent generation of the neflbE3 isoform. Importantly, over-expression of *neflbE4* ameliorated the impairment of motor neuron axonal length in TDP-43 KD mice, wherease *neflbE3* over-expression aggravated this phenotype. These observations, together with the overlapping motor and morphological deficits support interaction of TDP-43 and *nefl* in a common pathway, both biologically and pathogenetically.

### NEFL Splicing in human

In terms of mammalian NEFL, two possible splice variants are predicted for this transcript (Ensembl, ENSG00000277586) [78], [79]. This prediction is corroborated by the presence of multiple NEFL bands on Western blots of neuronal tissues, but they remain poorly characterized [80]. All together, the possible existence of a physiological or pathological splicing regulation of human NEFL and relevance to disease should be explored in light of the results of this study in zebrafish.

## Author Contributions

Experiments were performed by D-LD, M-LC and RMR. D-LD, and M-LC wrote the manuscript. The work was coordinated, revisited and approved by EK, HDD, and BJG.

## Acknowledgments

The authors would like to acknowledge Hortense de Calbiac for scientific insights, Anca Marian for technical assistance, Philippe Herbomel and Pierrick Jego for critical reading of the manuscript, as well as the dedicated platforms of Imagine and ICM Institutes for these studies.

