## Supplementary Figures for "Functional characterization of Neurofilament Light b splicing and *mis*balance in zebrafish"

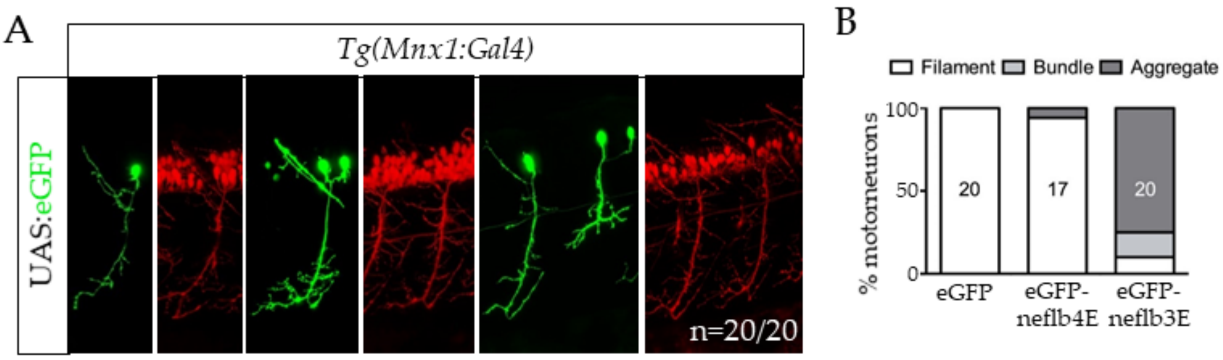


**Supplementary Figure 1. A,** *in vivo* observation of single motor neurons expressing eGFP from a UAS:eGFP plasmid injected at the 1-cell stage. Motor neurons expressing eGFP extend long and ramified axons. eGFP was detected in the whole cell, body, axon and ramifications, and did not induce silencing of the RFP reporter in any of the analyzed cells (n=20). **B**, quantification of percentages of filaments, bundles and aggregates structures developed in single motor neurons expressing eGFP-zNefl4E or eGFP-zNefl3E.

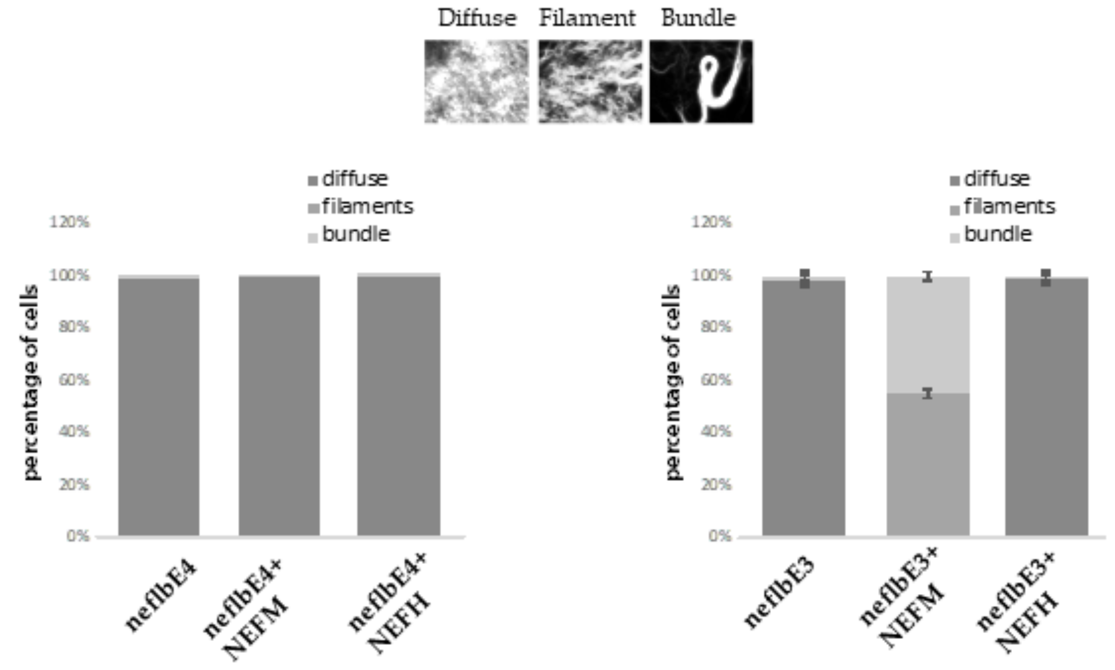


**Supplementary Figure 2.** Graphical representation of the percentage of cells carrying bundles, filaments and diffuse NFs structures in SW13$^{vim-}$ cells transfected with eGFP-neflbE4 (left) and eGFP-neflbE3 (right) alone or co-expressed with NEFM or mouse NEFH. The upper panel shows the typical shape of filaments, bundles or diffuse labeling that have been detect in transfected cells and quantified.

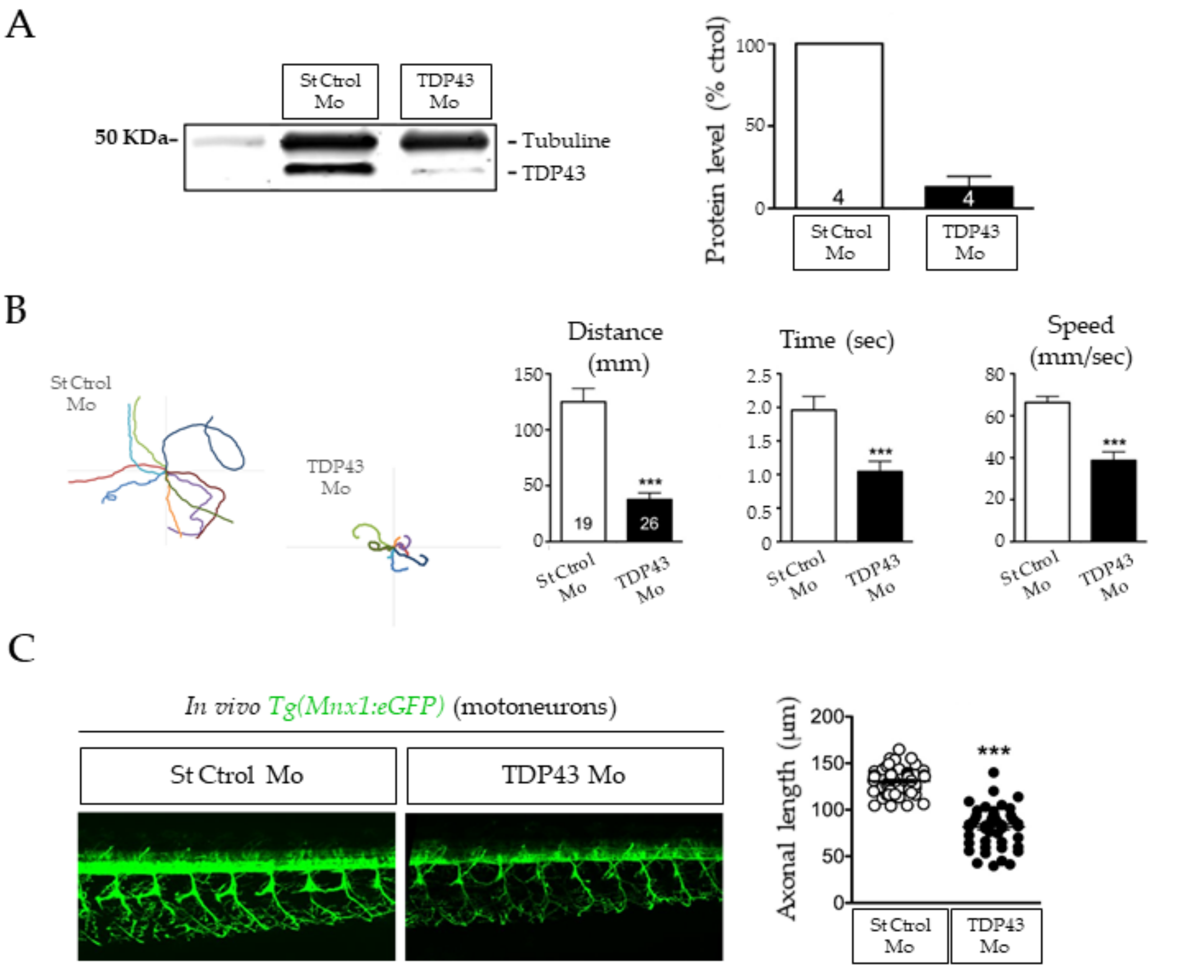


**Supplementary Figure 3: zTDP43 knock-down results in motor phenotype and axonal atrophy in zebrafish embryos. A**, TDP43 morpholino injection in zebrafish resulted in an almost 90% reduction of the TDP43 protein, as shown by WB. This KD lead to a strong and specific motor phenotype, as TDP43 morphants were unable to swim to the edges of a petri dish during TEER (**B**, left panel). Their displacement was shorter in distance, time and speed (B, right panel). **C**, motor axons of TDP43 morphants, revealed by GFP expression under the Mnx1 promoter *in vivo*, were shorter than control fish injected with ST Ctrol Mo. ***P<0.001.

**Supplementary Table 1**

| Gene of interest | Primer Forward | Primer Reverse |
|---|---|---|
| **bactin** | cccagacatcagggagtgat | tctctgttggctttgggatt |
| **RFP** | cgacatccccgactacttga | cttcttctgcattacggggc |
| **nefl** | gaggctgagaaggatgatgc | caagcattagggcacagtga |
| **nefl SV** | gaaggagaagaaacccgcttta | ctccttcttcaccaccttcc |
| **NEFL** | tgatggaagcgcgcaaag | ccccagcaccttcaactttc |

**Supplementary Movie 1 (related to Fig. 3). Somitic nerve fascicle growth in WT embryos.** *In vivo* time-lapse imaging of *Tg(Mnx1:eGFP)* zebrafish embryos injected with a control morpholino was performed during 10 hours starting from 16 hpf. *Mnx1:eGFP* line express GFP (green) in motor neurons. Axonal projections from motor neurons exit the spinal cord grouped as one nerve fascicle per somite, then grow along the somitic muscle all the way to its most ventral part while branching and connecting with muscle fibers.

**Supplementary Movie 2 (related to Fig. 3). Somitic nerve fascicle growth in NeflbSV embryos.** *In vivo* time-lapse imaging of *Tg(Mnx1:eGFP)* zebrafish embryos injected with the neflb SV morpholino was performed during 10 hours starting from 16 hpf. *Mnx1:eGFP* line express GFP (green) in motor neurons. Upon Neflb SV Mo injection, axonal projections initiate at the same developmental stage as in the controls, and they sprout in the right direction, ventrally towards the somitic muscles. However, motor axons constantly grew back and forth, and never reached normal length.
